# Interactions between environmental sensitivity and gut microbiota are associated with biomarkers of stress-related psychiatric symptoms

**DOI:** 10.1101/2023.03.30.534559

**Authors:** Shuhei Iimura, Satoshi Takasugi, Miyabi Yasuda, Yoshie Saito, Masashi Morifuji

**Affiliations:** Soka University; R&D Division, Meiji Co., Ltd

**Keywords:** environmental sensitivity, gut microbiota, brain-gut-microbiota axis, brain-gut axis, C-reactive protein, lipopolysaccharide-binding protein

## Abstract

**Background:** Humans vary in their sensitivity to stressful and supportive environments and experiences. Such individual differences in environmental sensitivity are associated with mechanisms of stress-related psychiatric symptoms. In recent years, researchers have focused on bidirectional interactions in the brain-gut-microbiota axis as a neurophysiological pathway contributing to the mechanisms of stress-related psychiatric symptoms, and evidence is rapidly accumulating.

**Methods:** Data on environmental sensitivity, gut microbiota, gut permeability (lipopolysaccharide-binding protein; LBP) and inflammation (C-reactive protein; CRP) were collected from 90 adults (50% female; *M*_age_ = 42.1; *SD*_age_ = 10.0). Environmental sensitivity was measured using a self-report questionnaire. Study participants’ feces were analyzed, and observed operational taxonomic units for richness, Shannon’s index for evenness, and phylogenetic diversity for biodiversity were evaluated as indicators of gut microbiota. In addition, participants’ serum was analyzed for CRP and LBP. We investigated whether the interaction between environmental sensitivity and gut microbiota is associated with biomarkers of inflammation and gut permeability.

**Results:** The interaction between environmental sensitivity and gut microbiota (excluding the Shannon’s index) explained the levels of these biomarkers. Individuals with high environmental sensitivity displayed higher levels of CRP and LBP, when the richness and diversity of the gut microbiota was low. However, even highly susceptible individuals had lower levels of CRP and LBP, when the richness and diversity of the gut microbiota was high.

**Conclusions:** Our study indicates that high environmental sensitivity can be a risk factor for inflammation and gut permeability, when the gut microbiota diversity is low, suggesting a brain-gut-microbiota axis interaction.

## Introduction

Environmental sensitivity, defined as individual differences in perception and processing of both positive and negative environments and stimuli (Pluess, 2015), moderates the relationship between environmental influences, including adversarial or supportive ones, and stress-related psychiatric symptoms, including depressive symptoms and anxiety, “for better *and* for worse” (Boyce & Ellis, 2005; Belsky & Pluess, 2009; Greven et al., 2019). Several genotypes (Keers et al., 2016), neurophysiological reactivity (Shakiba, Ellis, Bush & Boyce, 2020), and personality/temperamental factors (Aron et al., 2012) are associated with varying degrees of environmental sensitivity. Historically, high environmental sensitivity has been considered a risk/vulnerability factor for psychopathology in adversarial and stressful environments (Belsky & Pluess, 2009). However, current research on environmental sensitivity has redefined environmental sensitivity as a “plasticity” factor, rather than a “vulnerability” factor based on accumulating evidence (Belsky, Bakermans-Kranenburg & van Ijzendoorn, 2007; Belsky et al., 2009; Ellis, Boyce, Belsky, Bakermans-Kranenburg & van Ijzendoorn, 2011). Hence, individuals with high environmental sensitivity are not only more prone to increased psychopathology in stressful environments than those with low sensitivity (vulnerability), but also less prone to increased psychopathology if protective factors can be increased (vantage sensitivity). Indeed, several intervention studies have shown that individuals with high environmental sensitivity are more susceptible to the effects of resilience psychoeducation programs (Kibe, Suzuki, Hirano & Boniwell., 2020; Pluess & Boniwel, 2015) and cognitive-behavioral therapy (Keers et al., 2016) than those with low sensitivity, resulting in reported reductions in their depressive symptoms. Current research suggests that gut bacteria are a biological environmental factor that produce a variety of psychiatric symptoms through the brain-gut-microbiota axis (Zheng et al., 2016). In light of these recent observations, this study focuses on gut microbiota diversity, and examines how individual differences in environmental sensitivity are associated with biomarkers of stress-related psychiatric disorders.

Over the past decade, there has been a rapid accumulation of research findings on the bidirectional interaction between the brain and the gut microbiota (Mayer & Tillisch, 2011; Mayer, Savidge & Shulman, 2014), and these data have indicated the potential for these interactions to serve as new targets for the treatment of a wide range of psychiatric disorders, including depression and anxiety (Martin, Osadchiy, Kalani & Mayer, 2018; Winter, Hart, Charlesworth & Sharpley, 2018). Although the causal relationship has not yet been fully elucidated, the gut microbiota interacts with the brain through neuroimmune, neuroendocrine, and neural pathways, and thus, changes in the overall composition of the gut microbiota are associated with mood states (Winter et al., 2018). One pathway, suggested by several studies using animal models, is that stress exposure and the resulting depression-like behavioral changes in the brain affect gut microbiota diversity. For instance, it has been reported that rats subjected to an inevitable electric shock exhibit alterations in the composition of their gut microbiota, including *Lactobacillus*, when compared to rats that did not receive the shock or the control rats (Zhang et al., 2019). As an alternative pathway, it has been suggested that alterations in the gut microbiota may be involved in changes in depressive behavior. For example, transplantation of fecal microbiota collected from patients with major depressive disorder into germ-free mice resulted in observed behavioral changes related to depression and anxiety (Zheng et al., 2016). It has also been reported that depressive symptoms are associated with reduced richness and diversity of the gut microbiota from studies using oral gavage of fecal gut microbiota from depressed patients to microbiota-deficient rats (Kelly et al., 2016). These studies suggest that the brain, gut, and microbiota influence each other in reciprocal ways.

Inflammation can be a potent factor in the onset and maintenance processes of stress-related psychiatric disorders. A wide range of inflammatory biomarkers have been established for psychiatric disorders, including depressive symptoms and anxiety, such as C-reactive protein (CRP), gut permeability, tumor necrosis factor-alpha (TNF-α), and interleukins (IL) (Miller, 2020; Miller, Maletic & Raison, 2009). Patients with major depression have been found to have higher levels of inflammation (Raison, Capuron & Miller, 2006; Zorrilla et al., 2001). With regard to this mechanism, theories and review papers suggests that there are the bidirectional relationship between inflammatory biomarkers and psychiatric symptoms, including abnormalities in neurotransmitter metabolism (e.g., synthesis, release, and reuptake of monoamines) and neuroendocrine function (e.g., hyperactivity of the hypothalamic-pituitary-adrenal [HPA] axis with corticotropin-releasing hormone, adrenocorticotropic hormone and cortisol release), triggered by stressors (Maes et al, 2009; Schiepers, Wichers & Maes, 2005). There is growing evidence that inflammatory biomarkers interact with symptoms of stress-related psychiatric disorders. A longitudinal study showed that while the level of CRP at baseline predicted depressive symptoms surfacing approximately ten years later, depressive symptoms at baseline did not predict the level of CRP at ten years later, suggesting that inflammation precedes depressive symptoms (Gimeno et al., 2009).

Several meta-analyses have also reported that antidepressant treatment in patients with major depression reduces levels of CRP, IL-8 and TNF-α (Hiles, Baker, de Malmanche & Attia, 2012; Liu et al., 2020). As existing theory and findings suggest, the level of inflammation may be a key variable in understanding the mechanisms of stress-related psychiatric disorders and testing treatment efficacy.

Based on the discussion thus far, individual differences in environmental sensitivity, gut microbiota, and inflammatory biomarkers can correlate with each other. However, the relevance of these three variables, especially considering individual differences in environmental sensitivity, has not been examined to date. Given the established theory of environmental sensitivity (Boyce & Ellis, 2005; Belsky & Pluess, 2012, Pluess, 2015), higher environmental sensitivity may be associated with higher inflammation induced by stressors, but the relationship may be moderated by gut microbiota diversity. It is expected that the diversity of the gut microbiota would serve as a protective factor in highly susceptible individuals compared to those with lower sensitivity, and that a higher degree of diversity and richness of the microbiota would be less likely to increase the level of inflammatory indicators, biomarkers of stress-related psychiatric symptoms. This study examined this hypothetical relationship.

## Methods and Materials

### Study Procedure and Participants

We initially recruited 110 adults, who participated in the ongoing project “Identification of New Biomarkers for Intestinal Barrier Function,” and obtained consent to participate in this study from 90 (50% female; *M*_age_ = 42.1; *SD*_age_ = 10.0) of them. We asked them to respond to a web-based psychometric scale. The participants underwent a physical examination, blood analyses, sugar tolerance test, and stool test. Two years later, the participants responded to several psychological scales online. Stool test data were not available from two of the participants, so data from 88 participants were used in the main analysis related to gut microbiota. Detailed characteristics of the participants are shown in Supplemental Table S1 (https://bit.ly/3JAw0GX). The study design was preregistered at the University Hospital Medical Information Network (https://www.umin.ac.jp/english/) (Identifier: UMIN000047571). This study was approved by the Ethics Committee of Meiji Co., Ltd. (Tokyo, Japan) Institutional Review Board (No. 2021-012), considering the guidelines from the Declaration of Helsinki. All participants signed the informed consent form digitally.

### Personality Traits

#### Environmental Sensitivity

To measure personality traits associated with environmental sensitivity, we used the 10-item Japanese version of the Highly Sensitive Person Scale (Iimura et al., 2022). This scale measures susceptibility to both negative and positive environmental influences, and includes items such as “*Do changes in your life shake you up?*”, “*Are you easily overwhelmed by strong sensory input?*”, and “*Do you notice and enjoy delicate or fine scents, tastes, sounds, works of art?*”. All items are shown in Supplemental Table S2. Each item was rated on a 7-point Likert-type scale, ranging from 1 (*strongly disagree*) to 7 (*strongly agree*). Cronbach’s alpha, a measure of the scale’s internal consistency, was adequate at 0.85 (Omega total = .86). We used the 10-item mean for the analysis. It is generally supported that personality traits are psychological factors that remain stable within individuals over time (Costa & McCrae, 1988), and environmental sensitivity is no exception (Iimura, 2021).

### Biomarkers of Stress-Related Psychiatric Symptoms

#### C-Reactive Protein (CRP) and Lipopolysaccharide-Binding Protein (LBP)

Fasting venous blood was collected and serum separated by centrifugation. Serum samples were then stored at -80°C, and the serum levels of high-sensitivity CRP were measured by V-PLEX Vascular Injury Panel 2 Human Kit (Meso Scale Diagnostics, MD, US). CRP is reported to be an indicator that remains somewhat stable within individuals over time (Macy, Hayes & Tracy, 1997). Serum levels of LBP were measured using an LBP Human ELISA Kit (Hycult Biotech, Uden, The Netherlands). Based on previous studies (Gonzalez-Quintela et al., 2013; Taylor, Lehman, Kiefe & Seeman, 2006), both CRP and LBP were log-transformed and used in the analysis, due to their skewed distributions. The characteristics of these distributions are shown in Supplementary Figure S1.

### Gut Microbiota Diversity

#### Operational Taxonomic Units, Shannon’s Index, and Phylogenetic Diversity

Feces were collected by the participants themselves on any day during the experiment. Fecal samples were homogenized using FastPrep-24 5G (MP Biomedicals, Irvine, CA, US) with 0.1 mm Zirconia beads (EZ-Extract for DNA/RNA, AMR, Tokyo, Japan). Then, DNA was extracted from fecal samples using the QIAamp DNA Stool Mini Kit (QIAGEN, Hilden, Germany), according to “Protocol Q” (Costea et al., 2017), with a slight modification: A reduced time of beads beating step from 8 minutes to 4 minutes. The V3-V4 region of the 16S ribosomal RNA gene was amplified by PCR with universal bacterial primer sets (5’-TCGTCGGCAGCGTCAGATGTGTATAAGAGACAGCCTACGGGNGGCWGCAG-3’ and 5’-GTCTCGTGGGCTCGGAGATGTGTATAAGAGACAGGACTACHVGGGTATCTAATCC-3’) and was sequenced using MiSeq Reagent kit v3 (600 cycle) (Illumina Inc., California, US). The sequence data were analyzed using QIIME2 (https://qiime2.org/) to calculate alpha diversity. In this study, three indices were used to evaluate alpha diversity of gut microbiota (Nikolova et al., 2021): Observed operational taxonomic units (OTUs; observed species) for richness, Shannon’s index for evenness and richness/unevenness, and Faith’s phylogenetic diversity (Faith, 1992), which indicates biodiversity.

### Data Analysis

All the data in this study were analyzed using R version 4.2.2 (R Core Team, 2022). The R code used for the analysis has been uploaded to the Open Science Framework (https://bit.ly/3JAw0GX). The statistical significance level for the series of analyses was set at 5%.

#### Preliminary Analysis

Prior to the main analysis, means and standard deviations for each variable and bivariate correlation coefficients between each variable were calculated. Gender differences in each variable were also tested.

#### Analysis for Environmental Sensitivity × Gut Microbiota Diversity Interaction

The interaction effects of environmental sensitivity and gut microbiota diversity on the inflammatory biomarker (i.e., CRP) and gut permeability biomarker (i.e., LBP; Canipe et al., 2014; Stehle et al., 2012) were examined using hierarchical multiple regression analysis (Aiken & West, 1991). In Step 1, gender, age, BMI, environmental sensitivity, and gut microbiota diversity (observed OTUs, Shannon’s index, and phylogenetic diversity) were entered as independent variables, and the two biomarkers (i.e., CRP and LBP) involved in psychiatric symptoms were entered as dependent variables. In Step 2, an interaction term between environmental sensitivity and gut microbiota diversity was added to the independent variables entered in Step 1.

#### Analysis for Moderating Effect of Gut Microbiota Diversity

If the interaction term was significant in Step 2, a simple slope test was used to examine the moderating effect of gut microbiota diversity on the association between environmental sensitivity and biomarkers of psychopathology. Regression coefficients were then estimated by substituting *M*-1*SD*, Mean, and *M*+1*SD* for the gut microbiota diversity in the model, respectively (Aiken & West, 1991).

## Results

### Preliminary Analysis

Means and standard deviations for each variable, and the bivariate correlation coefficients between variables are shown in Table 1. No correlation was found between environmental sensitivity and biomarkers (i.e., CRP and LBP) of psychopathology, at least at the bivariate level. In addition, environmental sensitivity was not correlated with gut microbiota diversity (i.e., observed OTUs, Shannon’s index, and phylogenetic diversity). Observed OTUs showed a weak negative correlation with log-normalized CRP (*r* = -.23, *p* = .029). Correlations between the two biomarkers of stress-related symptoms were strong (*r* = .78, *p* <.001). Correlations between measures of gut microbiota diversity were also strong (*r* = 56 – .88, *p*s <.001), particularly between observed OTUs and phylogenetic diversity (*r* = .88, *p* <.001). Males (*M* = 7.70, *SD* = 1.66) showed higher mean log-normalized CRP than females (*M* = 6.96, *SD* = 1.08) (*t*(88) = 2.50, Cohen’s *d* = 0.52, *p* = .014). However, females showed higher mean levels of observed OTUs (male: *M* = 211.34, *SD* = 58.69, female: *M* = 267.98, *SD* = 78.83; *t*(86) = 3.82, *d* = 0.81, *p* <.001) and phylogenetic diversity (male: *M* = 14.24, *SD* = 3.21, female: *M* = 17.60, *SD* = 4.37; *t*(86) = 4.11, *d* = 0.87, *p* <.001) than males.

**Table 1.** Means, standard deviations, and correlations among variables

|  | *M* | *SD* | 1 | 2 | 3 | 4 | 5 | 6 | 7 | 8 |
| --- | --- | --- | --- | --- | --- | --- | --- | --- | --- | --- |
| 1. Gender | — | — | — |  |  |  |  |  |  |  |
| 2. Age | 42.10 | 10.05 | -.05 | — |  |  |  |  |  |  |
| 3. BMI | 22.75 | 3.40 | -.41** | .21 | — |  |  |  |  |  |
| 4. Sensitivity | 4.37 | 1.02 | .20 | .01 | -.07 | — |  |  |  |  |
| 5. CRP log | 7.33 | 1.44 | -.26* | -.02 | .54** | .15 | — |  |  |  |
| 6. LBP log | 2.58 | 0.26 | -.19 | .16 | .52** | .14 | .78** | — |  |  |
| 7. OTU | 239.66 | 74.73 | .38** | -.03 | -.32** | .18 | -.23* | -.17 | — |  |
| 8. Shannon | 5.65 | 0.55 | .11 | .13 | -.12 | .11 | -.09 | -.13 | .69** | — |
| 9. PD | 15.92 | 4.17 | .41** | -.09 | -.31** | .08 | -.15 | -.13 | .88** | .56** |

*Note.* BMI = Body Mass Index; Sensitivity = environmental sensitivity; CRP = C-reactive protein; LBP = lipopolysaccharide binding protein; OTU = observed operational taxonomic units (observes species); PD = phylogenetic diversity
\* $p < .05$ , \*\* $p < .01$

### Environmental Sensitivity **×** Gut Microbiota Diversity Interaction

#### Effect on CRP

As shown in Table 2, the interaction term between environmental sensitivity and observed OTUs was a significant predictor for CRP (*b* = -0.004, *SE* = 0.002, β = -.188, *p* = .039), even statistically controlling for the effects of other variables. The coefficient of determination, *R*^2^, increased from that in Step 1 when the interaction term between environmental sensitivity and observed OTUs was added in Step 2 (Δ*R*^2^ = .03, *F*[1, 81] = 4.41, *p* = .039). The coefficient of determination for the final regression model, Step 2, was *R*^2^ = 0.39 (*F*[6, 81] = 8.61, *p* <.001). The environmental sensitivity × Shannon interaction was not associated with CRP (Table 3; *b* = -0.485, *SE* = 0.254, β = -.171, *p* = .060), but the environmental sensitivity × phylogenetic diversity interaction predicted CRP (Table 4; *b* = -0.070, *SE* = 0.034, β = -.190, *p* = .040). Adding the environmental sensitivity × phylogenetic diversity interaction term increased the coefficient of determination (Δ*R*^2^ = .03, *F*[1, 81] = 34.33, *p* = .040). The coefficient of determination for this model (Step 2) was *R*^2^ = .39 (*F*[6, 81] = 8.46, *p* <.001).

**Table 2.**
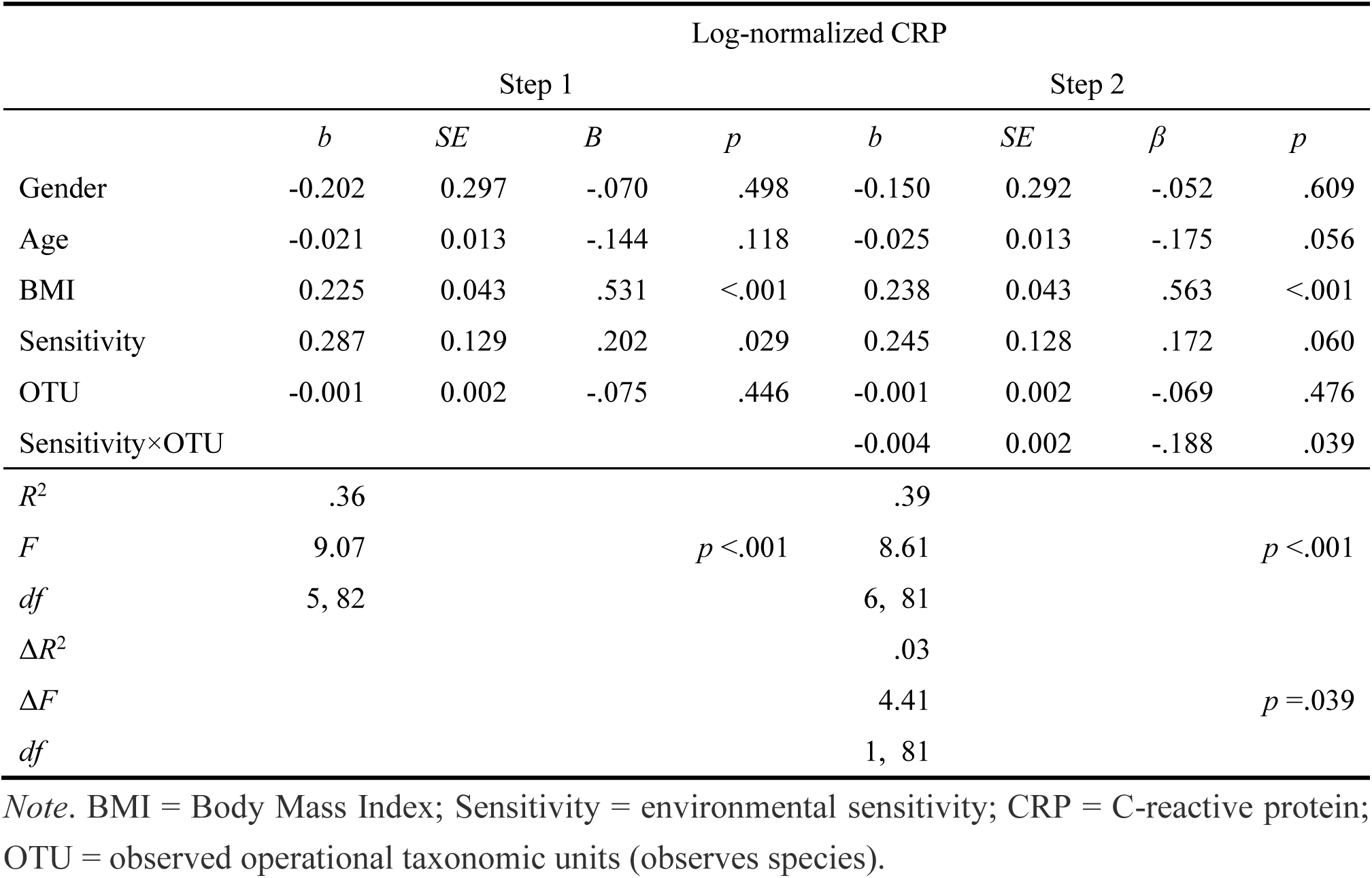
Interaction effects between environmental sensitivity and observed operational taxonomic units predicting C-reactive protein (*N* = 88)

**Table 3.**
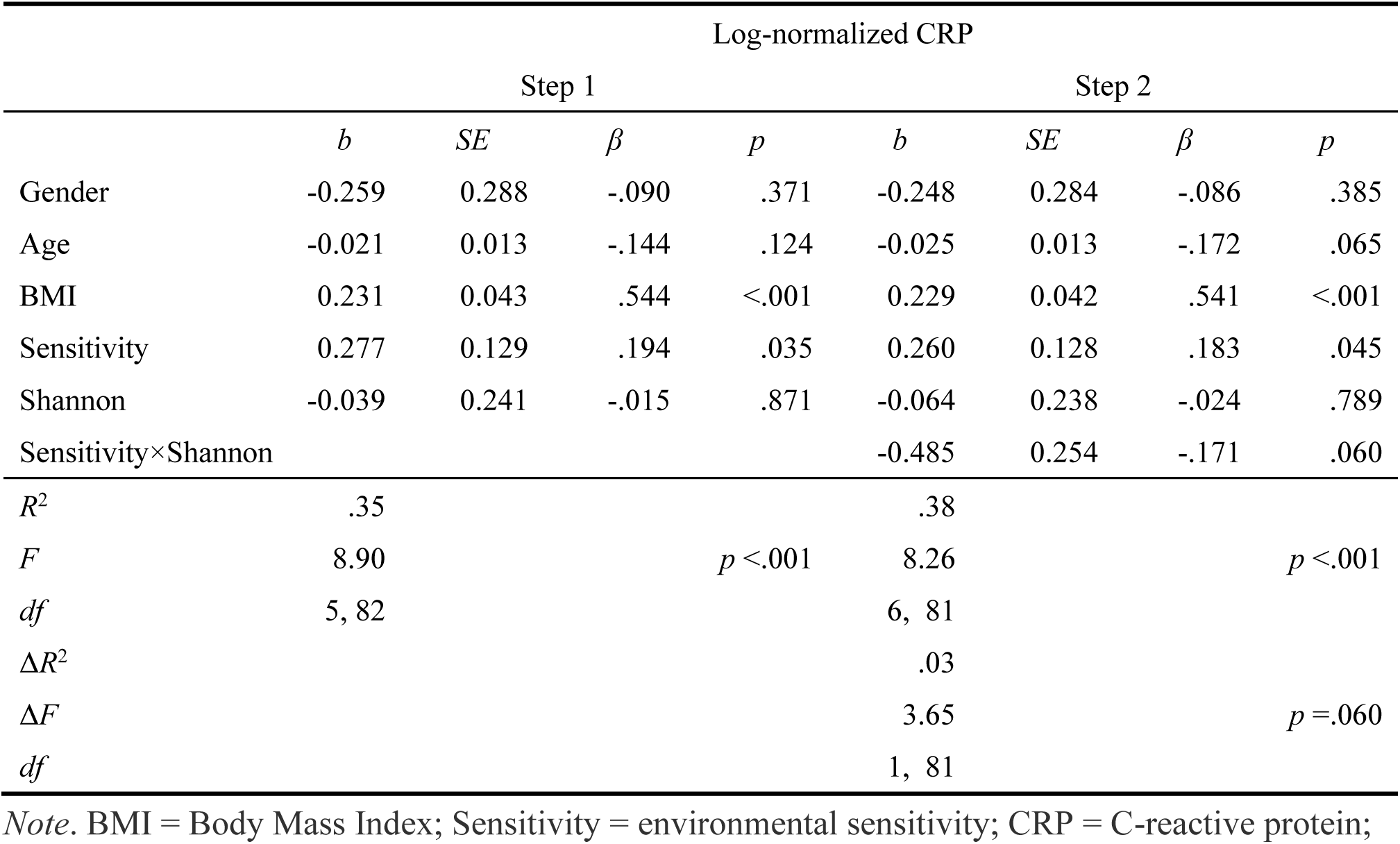
Interaction effects between environmental sensitivity and Shannon predicting C-reactive protein (*N* = 88)

**Table 4.** Interaction effects between environmental sensitivity and phylogenetic diversity predicting C-reactive protein (*N* = 88)

|  | Log-normalized CRP |  |  |  |  |  |  |  |
| --- | --- | --- | --- | --- | --- | --- | --- | --- |
|  | Step 1 |  |  |  | Step 2 |  |  |  |
| | *b* | *SE* | $\beta$ | *p* | *b* | *SE* | $\beta$ | *p* |
| Gender | -0.293 | 0.303 | -.102 | .336 | -0.238 | 0.298 | -.082 | .428 |
| Age | -0.021 | 0.013 | -.145 | .116 | -0.024 | 0.013 | -.166 | .070 |
| BMI | 0.234 | 0.043 | .552 | <.001 | 0.237 | 0.042 | .559 | <.001 |
| Sensitivity | 0.274 | 0.129 | .193 | .036 | 0.226 | 0.128 | .159 | .082 |
| PD | 0.012 | 0.034 | .033 | .739 | -0.003 | 0.034 | -.010 | .919 |
| Sensitivity×PD |  |  |  |  | -0.070 | 0.034 | -.190 | .040 |
| $R^2$ | .35 | | | | .39 | | | |
| *F* | 8.92 | | | $p < .001$ | 8.46 | | | $p < .001$ |
| *df* | 5, 82 |  |  |  | 6, 81 |  |  |  |
| $\Delta R^2$ | | | | | .03 | | | |
| $\Delta F$ | | | | | 34.33 | | | $p = .040$ |
| *df* |  |  |  |  | 1, 81 |  |  |  |

*Note.* BMI = Body Mass Index; Sensitivity = environmental sensitivity; CRP = C-reactive protein; PD = phylogenetic diversity

#### Effect on LBP

Similar results to CRP were obtained when LBP was the dependent variable. As shown in Table 5, the environmental sensitivity × observed OTUs interaction was associated with LBP (*b* = -0.001, *SE* <0.001, β = -.235, 95%CI [-0.419, -0.052], *p* = .013). Adding this interaction term in Step 2, *R*^2^ is increased from Step 1 (Δ*R*^2^ = .04, *F*[1, 81] = 6.50, *p* = .013). The *R*^2^ at Step 2 was .39 (*F*[6, 81] = 7.32, *p* <.001). The environmental sensitivity × Shannon interaction was not a predictor of LBP (Table 6; *b* = -0.060, *SE* = 0.047, β = -.119, *p* = .206). The interaction term between environmental sensitivity and phylogenetic diversity was associated with LBP (Table 7; *b* = -0.015, *SE* = 0.006, β = -.224, *p* = .019). When this interaction term was added in Step 2, *R*^2^ increased (Δ*R*^2^ = .05, *F*[1, 81] = 5.70, *p* = .019). *R*^2^ in Step 2 was .35 (*F*[6, 81] = 7.12, *p* <.001).

In summary, levels of inflammation were shown to be associated with the interaction between environmental sensitivity and gut microbiota diversity.

**Table 5.**
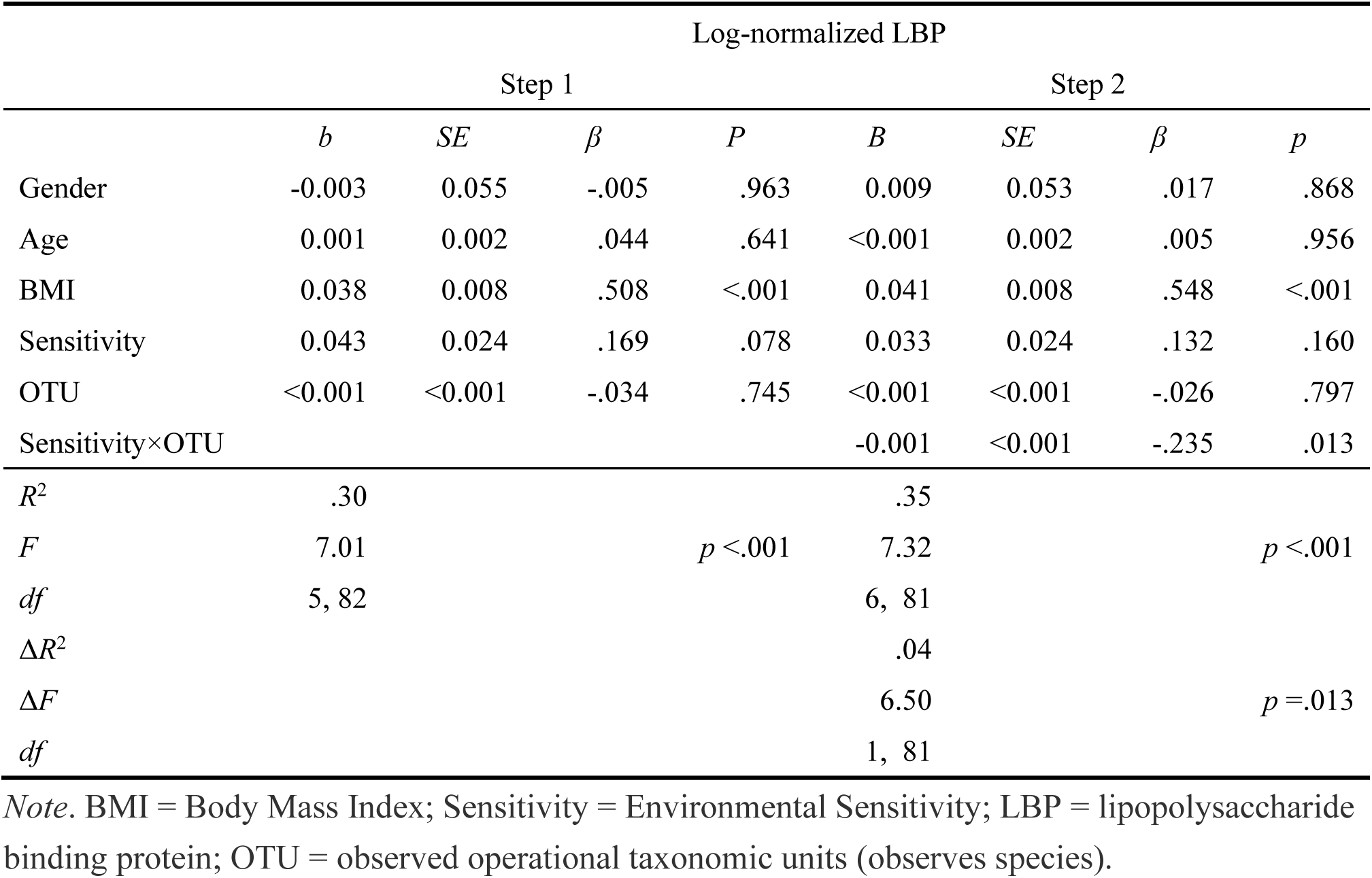
Interaction effects between environmental sensitivity and observed operational taxonomic units predicting lipopolysaccharide binding protein (*N* = 88)

**Table 6.**
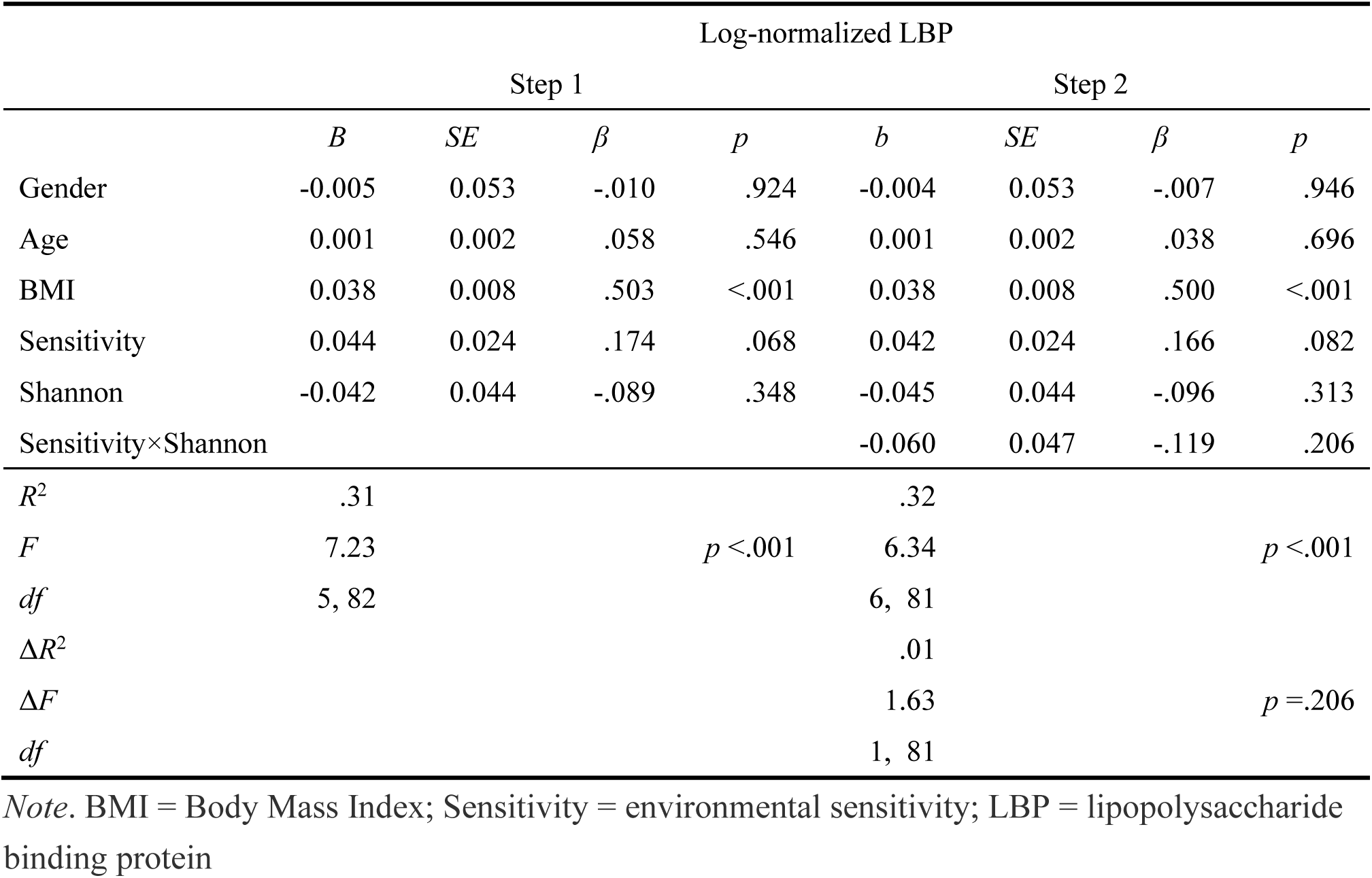
Interaction effects between environmental sensitivity and Shannon predicting lipopolysaccharide binding protein (*N* = 88)

**Table 7.**
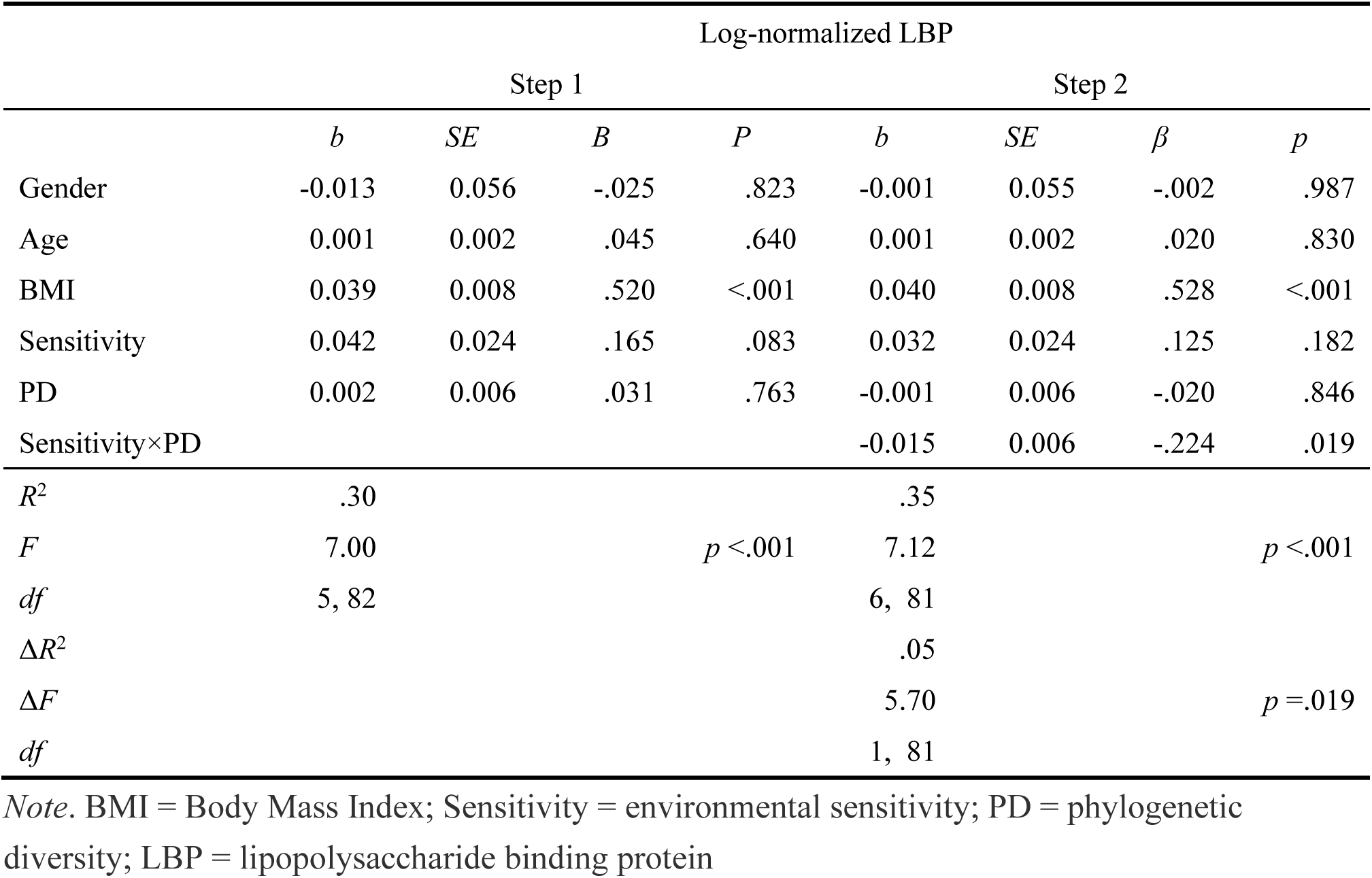
Interaction effects between environmental sensitivity and phylogenetic diversity predicting lipopolysaccharide binding protein (*N* = 88)

### Simple Slope Analysis

#### Effect on CRP

We performed simple slope tests on two indices of gut microbiota diversity (i.e., observed OTUs and phylogenetic diversity) for which we found an interaction effect (Figure 1). Individuals with high environmental sensitivity showed no association with CRP when observed OTUs were high (*M*+1*SD*; *b* = -.0.029, *SE* = 0.197, β = -.021, *p* = .882), but CRP was high when observed OTUs were low (*M*-1*SD*; *b* = .519., *SE* = 0.168, β = .365, *p* = .003) (Figure 1A). High environmental sensitivity was not associated with CRP when phylogenetic diversity was high (*M*+1*SD*; *b* = -.0.065, *SE* = 0.206, β = -.046, *p* = .752), but higher CRP when phylogenetic diversity was low (*M*-1*SD*; *b* = 0.518, *SE* = 0.172, β = .364, *p* = .003) (Figure 1B).

#### Effect on LBP

Similar interaction effects to CRP were also observed for LPB. Higher environmental sensitivity was not related to LBP when observed OTUs were high (*M*+1*SD*; *b* = -0.028, *SE* = 0.036, β = -.109, *p* = .446), but LBP was higher when observed OTUs were low (*M*-1*SD*; *b* = 0.094, *SE* = 0.031, β = .373, *p* = .003) (Figure 1C). Greater environmental sensitivity was not linked to LBP when phylogenetic diversity was high (*M*+1*SD*; *b* = -0.030, *SE* = 0.038, β = -.117, *p* = .435), but was linked to LBP when phylogenetic diversity was low (*M*-1*SD*; *b* = 0.093, *SE* = 0.032, β = .368, *p* = .004) (Figure 1D).

**Figure 1.**
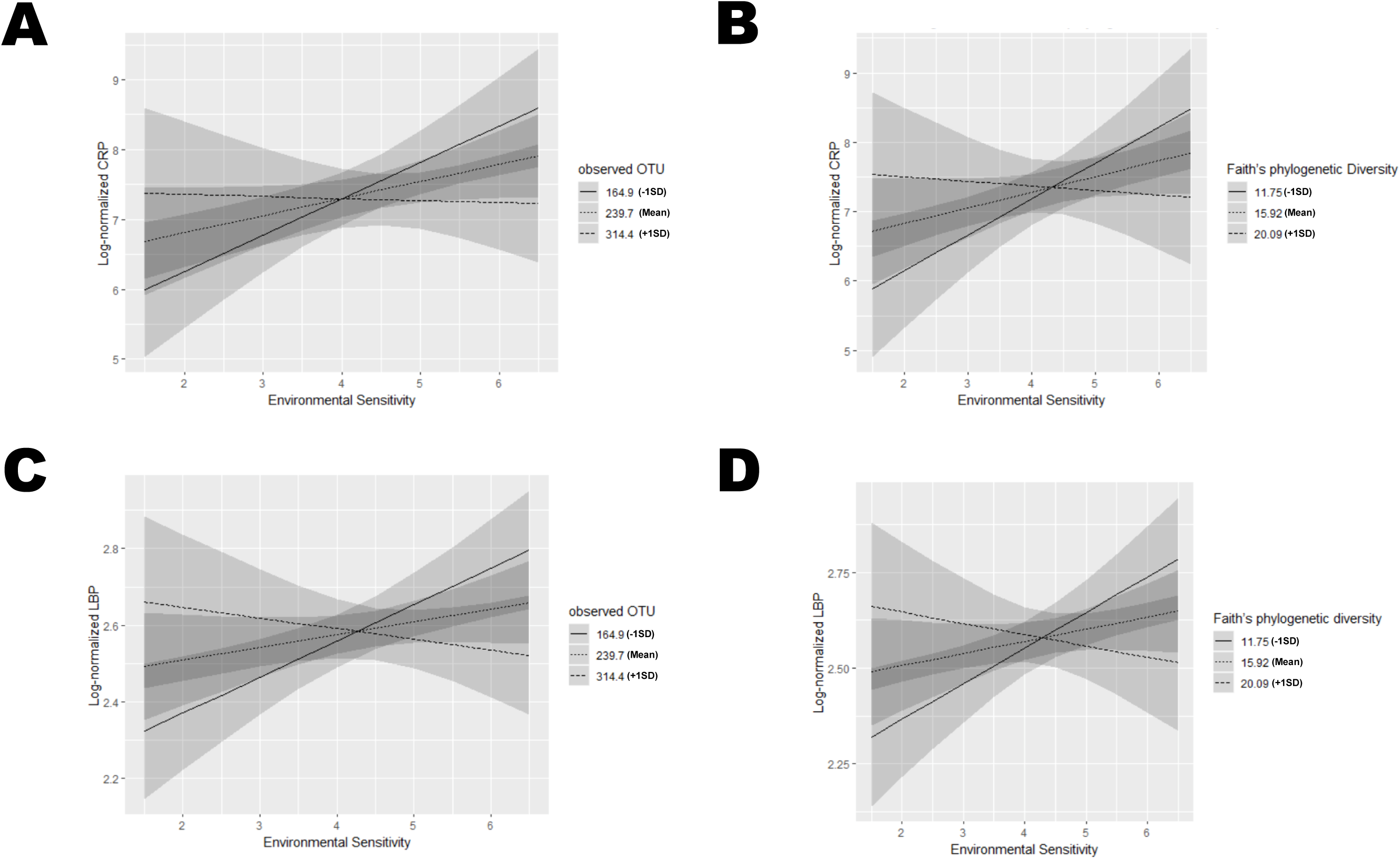
Moderating effects of environmental sensitivity gut microbiota diversity on C-reactive protein and lipopolysaccharide binding protein (*N* = 88). **(A)** Interaction effects between environmental sensitivity and observed operational taxonomic units on C-reactive protein. The three regression lines, from top to bottom, are for observed operational taxonomic units at *M*-1*SD* (= 164.9), Mean (= 239.7), and *M*+1*SD* (= 314.4), respectively. **(B)** Interaction effects between environmental sensitivity and phylogenetic diversity on C-reactive protein. The three regression lines, from top to bottom, are for phylogenetic diversity at *M*-1*SD* (= 11.8), Mean (= 15.9), and *M*+1*SD* (= 20.1), respectively. **(C)** Interaction effects between environmental sensitivity and observed operational taxonomic units on lipopolysaccharide binding protein. The three regression lines, from top to bottom, are for Alpha diversity at *M*-1*SD* (= 164.9), Mean (= 239.7), and *M*+1*SD* (= 314.4), respectively. **(D)** Interaction effects between environmental sensitivity and phylogenetic diversity on lipopolysaccharide binding protein. The three regression lines, from top to bottom, are for Alpha diversity at *M*-1*SD* (= 11.8), Mean (= 15.9), and *M*+1*SD* (= 20.1), respectively.

## Discussion

People differ in their susceptibility to external and internal environments and stimuli. The present study was the first to examine the association between gut microbiota and two biomarkers of stress-related psychiatric symptoms, considering individual differences in environmental sensitivity. We measured environmental sensitivity as a personality trait using a self-report questionnaire. Higher environmental sensitivity is not merely associated with greater vulnerability to negative influences from factors such as stressors, but also with greater benefit from protective factors such as a supportive environment and psychotherapy (Belsky & Pluess, 2012; Pluess, 2015; Greven et al., 2019). In this study, CRP and LBP were measured as indicators of biomarkers of psychopathology, which were used as outcome variables. It has been suggested that these higher degrees are associated with higher stress-related psychiatric symptoms, including depression and anxiety (Copeland et al., 2012; Gimeno et al., 2009; Iordache et al., 2022). To assess the function of the gut microbiota, we used observed OTUs for richness, Shannon’s index for evenness, and phylogenetic diversity for biodiversity, respectively, in our analysis. It has been previously reported that richness and diversity of the gut microbiota are associated with depressive symptoms (Kelly et al., 2016). We predicted that individual differences in environmental sensitivity would vary the extent to which the gut microbiota functions as a risk or protective factor, and consequently would show different associations with the levels of biomarkers of stress-related psychiatric symptoms.

Consequently, our data revealed that particularly highly sensitive individuals were not associated with higher levels of two biomarkers of psychopathology (i.e., CRP and LBP) if the richness and diversity of the microbiota were high; whereas, the level of CRP and LBP were higher when the richness and diversity of the microbiota was low.

This means that high environmental sensitivity is not necessarily a risk factor for stress-related psychopathology, suggesting that the richness and diversity of the gut microbiota may serve as a protective factor. Consistent with evolutionary developmental psychology theory and its findings (Belsky & Pluess, 2012; Ellis et al., 2011), our data suggest that high environmental sensitivity is not merely a “vulnerability” factor but a “plasticity” factor.

The present study showed for the first time that the association between individual differences in environmental sensitivity and two biomarkers of psychopathology (i.e., CRP and LBP) cannot be revealed by a simple bivariate correlation (see Table 1). In this study, we found that environmental sensitivity was positively associated with CRP and LBP by statistically controlling for BMI, gender, and other factors. Perhaps this indicates that the relationship between environmental sensitivity and CRP and LBP are not simple cause and effect relationships. It is plausible that individual differences in environmental sensitivity may lead to different levels of inflammatory responses, depending on susceptibility to risk factors (e.g., higher BMI and more stressors) and protective factors (e.g., good gut microbiota function and more social support). Studies examining the relationship between environmental sensitivity and self-reported depressive symptoms report that both are positively correlated (Liss, Mailloux & Erchull, 2008; Liss, Timmel, Baxley & Killingsworth, 2005). This represents the “dark side” of environmental sensitivity, i.e., the side in which individuals with high environmental sensitivity are more susceptible to adverse influence from stressful environments. Importantly, however, individuals with high environmental sensitivity experienced more positive life events during the week (Iimura, 2021), had better quality parent-child relationships (Rudolph et al., 2020), and took part in psychoeducational programs (Kibe et al, 2020; Nocentini, Menesini & Pluess, 2018; Pluess & Boniwel, 2015), have been reported to have better mental health on average, compared to less susceptible individuals. Considering the function of environmental sensitivity, comprehending the mechanisms responsible for the induction of stress-related psychiatric symptoms and associated inflammation necessitates contemplation of the interplay between internal and/or external environmental factors and environmental sensitivity.

A potential mechanism is that individual differences in environmental sensitivity would moderate the HPA axis activity to be more enhanced or more buffered when exposed to adverse environments (e.g., daily stressors) or supportive environments (e.g., high social support), consequently leading to different levels of inflammation, which impacts intestinal function and the composition of the gut microbiota (Al Bander, Nitert, Mousa & Naderpoor, 2020). This has been supported by a meta-analytic study, which reported that the serotonin transporter gene polymorphism (5-HTTLPR), one of the genetic markers of environmental sensitivity, is associated with cortisol responsiveness to acute psychosocial stress (Alexander et al., 2014; Miller, Wankerl, Stalder, Kirschbaum & Alexander, 2013), and individuals with the s-allele of 5HTTLPR have been suggested to be more susceptible to the environment than those with l-allele homozygotes (Belsky & Pluess, 2009; Van Ijzendoorn, Belsky & Bakermans-Kranenburg, 2012). Other possible mechanisms are that individuals with high environmental sensitivity may have a weakened intestinal barrier as a result of stressors. With low gut microbiota diversity, lipopolysaccharide (endotoxin) derived from bacteria enters the bloodstream through the intestinal tract, and increases inflammation (de Punder & Pruimboom, 2015). As a result, the level of CRP can increase.

Considering the effects of bacterial flora on the brain or related biomarkers based on these evidence, dietary interventions to increase the diversity of the gut microbiota and psychoeducational interventions to improve diet (Leeming, Johnson, Spector & Le Roy, 2019) may be more likely to benefit individuals who are more susceptible to higher environmental sensitivity. This is because previous studies have indicated that there are also individual differences in sensitivity to supportive environments and psychoeducational interventions (Iimura & Kibe, 2020; Kibe et al., 2020; Nocentini et al., 2018; Pluess & Boniwel, 2015).

This study has a number of limitations. The findings of this study rely on environmental sensitivity data collected via a questionnaire. Environmental sensitivity can also be measured by polygenic scores using genome-wide analysis (Keers et al., 2016), or by the degree of neurophysiological reactivity during stressful tasks (Weyn et al., 2022). Future studies should verify that the findings of this study can be replicated using these measures. In addition, the findings of this study were based on a correlational study design, and do not reveal a causal relationship between environmental sensitivity and gut microbiota and the brain. Future studies are expected to elucidate the reciprocal brain-gut-microbiota interaction through intervention or experimental manipulation of the gut microbiota. Finally, of the three indices of gut microbiota diversity, only Shannon’s index did not show a significant interaction with environmental sensitivity. Given this result, it is suggested that the number of species (i.e., observed OTUs and phylogenetic diversity) was a more important factor than homogeneity (i.e., Shannon’s index). Alternatively, it is possible that the statistical testing power was simply inadequate.

In conclusion, our study showed that interindividual differences between the two biomarkers of stress-related psychiatric symptoms were associated with an interaction between environmental sensitivity and gut microbiota diversity. Individuals with higher environmental sensitivity and lower levels of gut microbiota richness and diversity exhibited higher levels of inflammation and gut permeability. In contrast, even in highly susceptible individuals, a higher degree of richness and diversity of the microbiota did not increase the level of inflammation and gut permeability. These results suggest that high environmental sensitivity may be a risk factor for increased biomarkers of stress-related symptoms at lower levels of gut microbiota diversity. Moreover, individual differences in environmental sensitivity may be involved in brain-gut-microbiota interactions.

## References

Aiken, L. S., & West, S. G. (1991). Multiple regression: Testing and interpreting interactions. Sage Publications, Inc.

Al Bander, Z., Nitert, M. D., Mousa, A., & Naderpoor, N. (2020). The gut microbiota and inflammation: An overview. International Journal of Environmental Research and Public Health, 17(20). https://doi.org/10.3390/ijerph17207618

Alexander, N., Wankerl, M., Hennig, J., Miller, R., Zänkert, S., Steudte-Schmiedgen, S., Stalder, T., & Kirschbaum, C. (2014). DNA methylation profiles within the serotonin transporter gene moderate the association of 5-HTTLPR and cortisol stress reactivity. Translational Psychiatry, 4, e443. https://doi.org/10.1038/tp.2014.88

Belsky, J., Bakermans-Kranenburg, M. J., & van IJzendoorn, M. H. (2007). For better and for worse: Differential susceptibility to environmental influences. Current Directions in Psychological Science, 16, 300–304. https://doi.org/10.1111/j.1467-8721.2007.00525.x

Belsky, J., Jonassaint, C., Pluess, M., Stanton, M., Brummett, B., & Williams, R. (2009). Vulnerability genes or plasticity genes? Molecular Psychiatry, 14, 746–754. https://doi.org/10.1038/mp.2009.44

Belsky, J., & Pluess, M. (2009). Beyond diathesis stress: Differential susceptibility to environmental influences. Psychological Bulletin, 135, 885–908. https://doi.org/10.1037/a0017376

Boyce, W. T., & Ellis, B. J. (2005). Biological sensitivity to context: I. An evolutionary-developmental theory of the origins and functions of stress reactivity. Development and Psychopathology, 17, 271–301. https://doi.org/10.1017/s0954579405050145

Canipe, A., Chidumayo, T., Blevins, M., Bestawros, M., Bala, J., Kelly, P., … & Koethe, J. R. (2014). A 12 week longitudinal study of microbial translocation and systemic inflammation in undernourished HIV-infected Zambians initiating antiretroviral therapy. BMC Infectious Diseases, 14, 1–9. https://doi.org/10.1186/1471-2334-14-521

Copeland, W. E., Shanahan, L., Worthman, C., Angold, A., & Costello, E. J. (2012). Generalized anxiety and C-reactive protein levels: A prospective, longitudinal analysis. Psychological Medicine, 42, 2641–2650. https://doi.org/10.1017/S0033291712000554

Costa, P. T., & McCrae, R. R. (1988). Personality in adulthood: A six-year longitudinal study of self-reports and spouse ratings on the NEO Personality Inventory. Journal of Personality and Social Psychology, 54, 853–863.

Costea, P. I., Zeller, G., Sunagawa, S., Pelletier, E., Alberti, A., Levenez, F., … & Bork, P. (2017). Towards standards for human fecal sample processing in metagenomic studies. Nature Biotechnology, 35, 1069–1076. https://doi.org/10.1038/nbt.3960

de Punder, K., & Pruimboom, L. (2015). Stress induces endotoxemia and low-grade inflammation by increasing barrier permeability. Frontiers in Immunology, 6, 223. https://doi.org/10.3389/fimmu.2015.00223

de Villiers, B., Lionetti, F., & Pluess, M. (2018). Vantage sensitivity: A framework for individual differences in response to psychological intervention. Social Psychiatry and Psychiatric Epidemiology, 53, 545–554. https://doi.org/10.1007/s00127-017-1471-0

Ellis, B. J., Boyce, W. T., Belsky, J., Bakermans-Kranenburg, M. J., & Van Ijzendoorn, M. H. (2011). Differential susceptibility to the environment: An evolutionary-neurodevelopmental theory. Development and Psychopathology, 23, 7–28. https://doi.org/10.1017/S0954579410000611

Faith, D. P. (1992). Conservation evaluation and phylogenetic diversity. Biological Conservation, 61, 1–10. https://doi.org/10.1016/0006-3207(92)91201-3

Gimeno, D., Kivimäki, M., Brunner, E. J., Elovainio, M., De Vogli, R., Steptoe, A., Kumari, M., Lowe, G. D. O., Rumley, A., Marmot, M. G., & Ferrie, J. E. (2009). Associations of C-reactive protein and interleukin-6 with cognitive symptoms of depression: 12-year follow-up of the Whitehall II study. Psychological Medicine, 39, 413–423. https://doi.org/10.1017/S0033291708003723

Gonzalez-Quintela, A., Alonso, M., Campos, J., Vizcaino, L., Loidi, L., & Gude, F. (2013). Determinants of serum concentrations of lipopolysaccharide-binding protein (LBP) in the adult population: The role of obesity. PloS one, 8, e54600. https://doi.org/10.1371/journal.pone.0054600

Greven, C. U., Lionetti, F., Booth, C., Aron, E. N., Fox, E., Schendan, H. E., Homberg, J. (2019). Sensory processing sensitivity in the context of environmental sensitivity: A critical review and development of research agenda. Neuroscience & Biobehavioral Reviews, 8, 287–305. https://doi.org/10.1016/j.neubiorev.2019.01.009

Hiles, S. A., Baker, A. L., de Malmanche, T., & Attia, J. (2012). Interleukin-6, C-reactive protein and interleukin-10 after antidepressant treatment in people with depression: a meta-analysis. Psychological Medicine, 42, 2015–2026. https://doi.org/10.1017/S0033291712000128

Iimura, S. (2021). Highly sensitive adolescents: The relationship between weekly life events and weekly socioemotional well-being. British Journal of Psychology, 112, 1103–1129. https://doi.org/10.1111/bjop.12505

Iimura, S. (2022). Sensory-processing sensitivity and COVID-19 stress in a young population: The mediating role of resilience. Personality and Individual Differences, 184. https://doi.org/10.1016/j.paid.2021.111183

Iimura, S., Yano, K., & Ishii, Y. (2021). Environmental sensitivity in adults: Psychometric properties of the Japanese Version of the Highly Sensitive Person Scale 10-Item Version. Journal of Personality Assessment. https://doi.org/10.31234/osf.io/2h8jt

Iordache, M. M., Tocia, C., Aschie, M., Dumitru, A., Manea, M., Cozaru, G. C., Petcu, L., et al. (2022). Intestinal permeability and depression in patients with inflammatory bowel disease. Journal of Clinical Medicine, 11, 5121. http://dx.doi.org/10.3390/jcm11175121

Keers, R., Coleman, J. R. I., Lester, K. J., Roberts, S., Breen, G., Thastum, M., et al. (2016). A genome-wide test of the differential susceptibility hypothesis reveals a genetic predictor of differential response to psychological treatments for child anxiety disorders. Psychotherapy and Psychosomatics, 85, 146–158. https://doi.org/10.1159/000444023

Kelly, J. R., Borre, Y., O’ Brien, C., Patterson, E., El Aidy, S., Deane, J., et al. (2016). Transferring the blues: Depression-associated gut microbiota induces neurobehavioural changes in the rat. Journal of Psychiatric Research, 82, 109–118. https://doi.org/10.1016/j.jpsychires.2016.07.019

Kibe, C., Suzuki, M., Hirano, M., & Boniwell, I. (2020). Sensory processing sensitivity and culturally modified resilience education: Differential susceptibility in Japanese adolescents. PloS One, 15, 1–17. https://doi.org/10.1371/journal.pone.0239002

Leeming, E. R., Johnson, A. J., Spector, T. D., & Le Roy, C. I. (2019). Effect of diet on the gut microbiota: Rethinking intervention duration. Nutrients, 11. https://doi.org/10.3390/nu11122862

Liss, M., Mailloux, J., & Erchull, M. J. (2008). The relationships between sensory processing sensitivity, alexithymia, autism, depression, and anxiety. Personality and Individual Differences, 45, 255–259. https://doi.org/10.1016/j.paid.2008.04.009

Liss, M., Timmel, L., Baxley, K., & Killingsworth, P. (2005). Sensory processing sensitivity and its relation to parental bonding, anxiety, and depression. Personality and Individual Differences, 39, 1429–1439. https://doi.org/10.1016/j.paid.2005.05.007

Liu, J. J., Wei, Y. B., Strawbridge, R., Bao, Y., Chang, S., Shi, L., et al. (2020). Peripheral cytokine levels and response to antidepressant treatment in depression: A systematic review and meta-analysis. Molecular Psychiatry, 25, 339–350. https://doi.org/10.1038/s41380-019-0474-5

Macy, E. M., Hayes, T. E., & Tracy, R. P. (1997). Variability in the measurement of C-reactive protein in healthy subjects: implications for reference intervals and epidemiological applications. Clinical Chemistry, 43, 52–58.

Maes, M., Yirmyia, R., Noraberg, J., Brene, S., Hibbeln, J., Perini, G., et al. (2009). The inflammatory & neurodegenerative (I&ND) hypothesis of depression: Leads for future research and new drug developments in depression. Metabolic Brain Disease, 24, 27–53. https://doi.org/10.1007/s11011-008-9118-1

Martin, C. R., Osadchiy, V., Kalani, A., & Mayer, E. A. (2018). The brain-gut-microbiome axis. Cellular and Molecular Gastroenterology and Hepatology, 6, 133–148. https://doi.org/10.1016/j.jcmgh.2018.04.003

Mayer, E. A., Savidge, T., & Shulman, R. J. (2014). Brain-gut microbiome interactions and functional bowel disorders. Gastroenterology, 146, 1500–1512. https://doi.org/10.1053/j.gastro.2014.02.037

Mayer, E. A., & Tillisch, K. (2011). The brain-gut axis in abdominal pain syndromes. Annual Review of Medicine, 62, 381–396. https://doi.org/10.1146/annurev-med-012309-103958

Miller, A. H. (2020). Beyond depression: The expanding role of inflammation in psychiatric disorders. World Psychiatry, 19, 108. https://doi.org/10.1002/wps.20723

Miller, A. H., Maletic, V., & Raison, C. L. (2009). Inflammation and its discontents: The role of cytokines in the pathophysiology of major depression. Biological Psychiatry, 65, 732–741. https://doi.org/10.1016/j.biopsych.2008.11.029

Miller, R., Wankerl, M., Stalder, T., Kirschbaum, C., & Alexander, N. (2013). The serotonin transporter gene-linked polymorphic region (5-HTTLPR) and cortisol stress reactivity: A meta-analysis. Molecular Psychiatry, 18, 1018–1024. https://doi.org/10.1038/mp.2012.124

Nikolova, V. L., Hall, M. R., Hall, L. J., Cleare, A. J., Stone, J. M., & Young, A. H. (2021). Perturbations in gut microbiota composition in psychiatric disorders: A review and meta-analysis. JAMA Psychiatry, 78, 1343–1354. https://doi.org/10.1001/jamapsychiatry.2021.2573

Nocentini, A., Menesini, E., & Pluess, M. (2018). The personality trait of environmental sensitivity predicts children’s positive response to school-based antibullying intervention. Clinical Psychological Science, 6, 848–859. https://doi.org/10.1177/2167702618782194

Pluess, M. (2017). Vantage sensitivity: Environmental sensitivity to positive experiences as a function of genetic differences. Journal of Personality, 85, 38–50. https://doi.org/10.1111/jopy.12218

Pluess, M., & Belsky, J. (2013). Vantage sensitivity: Individual differences in response to positive experiences. Psychological Bulletin, 139, 901–916. https://doi.org/10.1037/a0030196

Pluess, M., & Boniwell, I. (2015). Sensory-processing sensitivity predicts treatment response to a school-based depression prevention program: Evidence of vantage sensitivity. Personality and Individual Differences, 82, 40–45. https://doi.org/10.1016/j.paid.2015.03.011

Raison, C. L., Capuron, L., & Miller, A. H. (2006). Cytokines sing the blues: Inflammation and the pathogenesis of depression. Trends in Immunology, 27, 24–31. https://doi.org/10.1016/j.it.2005.11.006

R Core Team (2022). R: A language and environment for statistical computing. R Foundation for Statistical Computing, Vienna, Austria. Retrieved from https://www.R-project.org/.

Rudolph, K. D., Davis, M. M., Modi, H. H., Fowler, C., Kim, Y., & Telzer, E. H. (2020). Differential susceptibility to parenting in adolescent girls: Moderation by neural sensitivity to social cues. Journal of Research on Adolescence, 30, 177–191. https://doi.org/10.1111/jora.12458

Schiepers, O. J., Wichers, M. C., & Maes, M. (2005). Cytokines and major depression. Progress in Neuro-Psychopharmacology and Biological Psychiatry, 29, 201–217. https://doi.org/10.1016/j.pnpbp.2004.11.003

Shakiba, N., Ellis, B. J., Bush, N. R., & Boyce, W. T. (2020). Biological sensitivity to context: A test of the hypothesized U-shaped relation between early adversity and stress responsivity. Development and Psychopathology, 32, 641–660. https://doi.org/10.1017/S0954579419000518

Stehle Jr, J. R., Leng, X., Kitzman, D. W., Nicklas, B. J., Kritchevsky, S. B., & High, K. P. (2012). Lipopolysaccharide-binding protein, a surrogate marker of microbial translocation, is associated with physical function in healthy older adults. Journals of Gerontology: Series A, 67, 1212–1218. https://doi.org/10.1093/gerona/gls178

Taylor, S. E., Lehman, B. J., Kiefe, C. I., & Seeman, T. E. (2006). Relationship of early life stress and psychological functioning to adult C-reactive protein in the coronary artery risk development in young adults study. Biological Psychiatry, 60, 819–824. https://doi.org/10.1016/j.biopsych.2006.03.016

Van Ijzendoorn, M. H., Belsky, J., & Bakermans-Kranenburg, M. J. (2012). Serotonin transporter genotype 5HTTLPR as a marker of differential susceptibility: A meta-analysis of child and adolescent gene-by-environment studies. Translational Psychiatry, 2, e147–6. https://doi.org/10.1038/tp.2012.73

Weyn, S., Van Leeuwen, K., Pluess, M., Goossens, L., Claes, S., Bosmans, G., et al. (2022). Individual differences in environmental sensitivity at physiological and phenotypic level: Two sides of the same coin? International Journal of Psychophysiology, 176, 36–53. https://doi.org/10.1016/j.ijpsycho.2022.02.010

Winter, G., Hart, R. A., Charlesworth, R. P. G., & Sharpley, C. F. (2018). Gut microbiome and depression: What we know and what we need to know. Reviews in the Neurosciences, 29, 629–643. https://doi.org/10.1515/revneuro-2017-0072

Zhang, K., Fujita, Y., Chang, L., Qu, Y., Pu, Y., Wang, S., Shirayama, Y., & Hashimoto, K. (2019). Abnormal composition of gut microbiota is associated with resilience versus susceptibility to inescapable electric stress. Translational Psychiatry, 9, 231. https://doi.org/10.1038/s41398-019-0571-x

Zheng, P., Zeng, B., Zhou, C., Liu, M., Fang, Z., Xu, X., et al. (2016). Gut microbiome remodeling induces depressive-like behaviors through a pathway mediated by the host’s metabolism. Molecular Psychiatry, 21, 786–796. https://doi.org/10.1038/mp.2016.44

Zorrilla, E. P., Luborsky, L., McKay, J. R., Rosenthal, R., Houldin, A., Tax, A., et al. (2001). The relationship of depression and stressors to immunological assays: A meta-analytic review. Brain, Behavior, and Immunity, 15, 199–226. https://doi.org/10.1006/brbi.2000.0597

