## Supplemental File for "Interactions between environmental sensitivity and gut microbiota are associated with biomarkers of stress-related psychiatric symptoms"

**----- Table of Contents -----**

**Table S1**: Detailed characteristics of study participants

Table S2: The Japanese version of the Highly Person Scale-10 item version

**Figure S1**: Histograms of key variables

**Table S1.** Detailed characteristics of study participants.

A: Number of people living together

| | *N* |
|---|---|
| One | 26 |
| Two | 18 |
| Three | 22 |
| Four | 17 |
| Five | 7 |
| Six | 0 |
| Seven or more | 0 |

B: Marital status

| | *N* |
|---|---|
| Married | 47 |
| Not married | 38 |
| Divorced, separated, or bereaved | 5 |

C: Number of sons or daughters

| | *N* |
|---|---|
| One | 15 |
| Two | 15 |
| Three | 6 |
| Four or more | 1 |

| | |
|---|---|
| None | 53 |

D: Annual family income

| | *N* |
|---|---|
| Less than 2 million yen | 7 |
| 2 million to 4 million yen | 18 |
| 4.01 million to 6 million yen | 17 |
| 6.01 million to 8 million yen | 22 |
| 8.01 million to 10 million yen | 12 |
| 1,001,000 to 12,000,000 yen | 6 |
| 12.01 million yen and above | 8 |

E: Educational background

| | *N* |
|---|---|
| Elementary and Junior High School | 0 |
| High School | 15 |
| Technical College/Junior College | 30 |
| Universities | 43 |
| Graduate Schools | 2 |

Table S2. The Japanese version of the Highly Person Scale-10 item version

| |
|---|
| Do changes in your life shake you up? |
| Are you easily overwhelmed by strong sensory input? |
| Do other people's moods affect you? |
| Do you get rattled when you have a lot to do in a short amount of time? |
| When you must compete or be observed while performing a task, do you become so nervous or shaky that you do much worse than you would otherwise? |
| Are you bothered by intense stimuli, like loud noises or chaotic scenes? |
| Are you made uncomfortable by loud noises? |
| Are you easily overwhelmed by things like bright lights, strong smells, coarse fabrics, or sirens close by? |
| Do you notice and enjoy delicate or fine scents, tastes, sounds, works of art? |
| Are you deeply moved by the arts or music? |

*Note*. Each item was rated on a 7-point Likert-type scale, ranging from 1 (strongly disagree) to 7 (strongly agree).

**Figure S1.** *Histograms of key variables.* (A) Environmental sensitivity. (B) Log-normalized C-reactive protein. (C) Log-normalized lipopolysaccharide binding protein. (D) Observed operational taxonomic units. (E) Shannon. (F) Faith’s phylogenetic diversity.

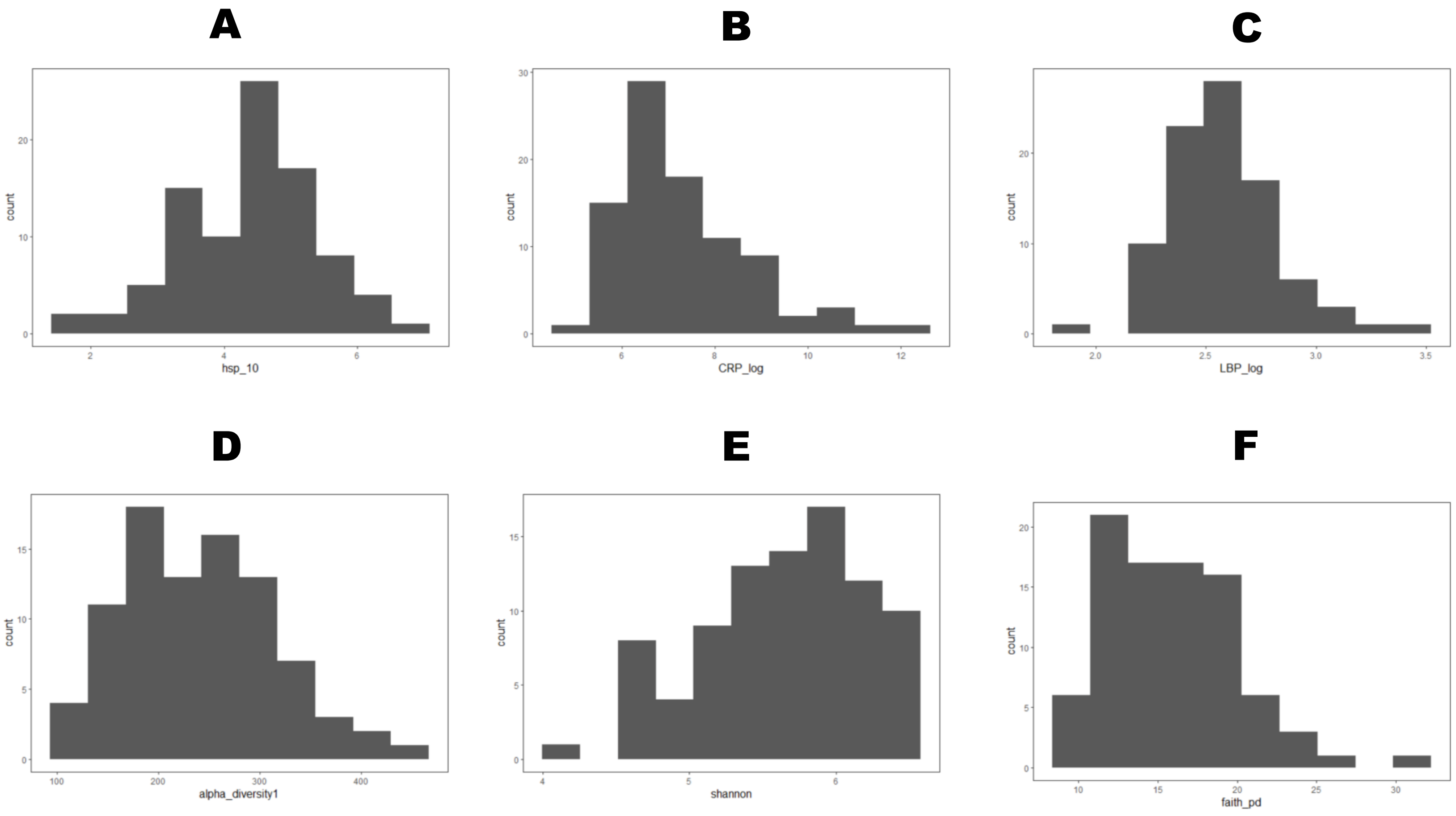
